# A surprising lack of presaccadic benefits during visual change detection

**DOI:** 10.1101/2023.01.01.522284

**Authors:** Priyanka Gupta, Devarajan Sridharan

## Abstract

Even before the eyes move, visual sensitivity improves at the target of the planned eye movement. Yet, it is unknown if such “presaccadic” benefits are merely sensory, or also influence decisional processes. We teased apart these contributions with signal detection theory, and discovered a surprising absence of presaccadic benefits in visual change detection tasks. Participants planned and executed saccades while concurrently detecting and localizing either orientation or contrast changes. Spatial choice bias reliably improved presaccadically but, surprisingly, without a concomitant increase in perceptual sensitivity. Additional investigation with an orientation estimation task, and a Bayesian “variable precision” model, revealed that sensory precision increased at the saccade target, but only for the most recent of two successive stimuli. Moreover, the recent stimulus perceptually biased feature estimates of the prior stimulus, rendering accurate change detection even more challenging. Our results uncover novel perceptual and decisional mechanisms that mediate presaccadic change detection.

**Lay Summary:** Planning a rapid eye movement (saccade) changes how we perceive our visual world. Even before we move the eyes visual sensitivity improves at the impending target of eye movements, a phenomenon termed “presaccadic attention”. We report a surprising lack of such presaccadic attention benefits in a common, everyday setting: change detection. In fact, presaccadic attention renders change detection more challenging by biasing percepts toward the most recent of successive stimuli at the saccade target location. With a Bayesian model, we show how such perceptual and choice biases induced by eye movement planning affect change detection behavior. Our findings may have critical implications for real-world scenarios, like driving, that involve rapid gaze shifts in dynamically changing environments.

## Introduction

Non-uniform visual acuity in the retina prevents perceiving the entire visual scene at high resolution all at once. Saccades are ballistic eye movements that enable us to bring into focus specific, important parts of the visual scene, in quick succession, for foveal analysis. But even before the eyes move visual sensitivity improves at the location of the impending saccade: a phenomenon termed “presaccadic attention”.

An extensive literature has characterized behavioral benefits of presaccadic attention. Both visual discrimination and identification accuracy have been reported to improve presaccadically^1–5^. In particular, many studies have reported presaccadic improvements in orientation discrimination accuracy^6–12^. Several mechanisms have been proposed to explain these effects. For example, behavioral evidence suggests that presaccadic attention narrows orientation tuning^13–15^, and preferentially enhances high spatial frequency information^13,16^ as well as the encoding of specific orientations^17^ thereby increasing discrimination accuracy for certain types of stimuli. Moreover, the orientation information content at the saccade target has been shown to bias processing both at the fovea^18^, and the peripheral visual field^19^, suggesting a selection in feature-space following spatial selection. In addition, saccade preparation increases the perceived contrast at the saccade target location^6^, and produces response gain-like effects^10^, suggesting a potential presaccadic gain in encoding strength. While the spatial spread of presaccadic attention is strongly determined by visual context^20,21^, improved discrimination at the saccade target location is accompanied by a decrement at other locations, suggesting a capacity-limited presaccadic selection mechanism^8,22^.

Despite this significant literature characterizing the effects of presaccadic attention, it is unclear how selection at the saccade target influences perceptual versus decisional processing^8,13^. The vast majority of previous literature relied on discrimination tasks employing conventional forced choice responses^5,6,9,12,20,23^; therefore, the generalizability of these findings regarding the mechanisms of presaccadic attention is limited. Importantly, these designs do not permit disambiguating sensory (sensitivity) from downstream decisional (criterion) effects.

Here, we employ signal detection theory and Bayesian modeling of behavior to distinguish perceptual and decisional mechanisms by which presaccadic attention facilitates visual change detection. In a surprising counterpoint to previous literature, we show that – despite producing sensory and decisional advantages at the saccade target location – presaccadic attention, paradoxically, does not benefit change detection. We developed a dual task paradigm to quantify orientation change detection ability, and with a recent signal detection theory model^24,25^, quantified distinct components of presaccadic attention – sensitivity and criteria – at the saccade target and other (non-saccade target) locations. Surprisingly, change detection sensitivity was not enhanced presaccadically, at the saccade target location. Rather, choice criterion was the lowest, and spatial choice bias highest, for reporting a change at the saccade target location. To further investigate these findings, we conducted another experiment where participants estimated, presaccadically, the orientation of one of two stimuli, presented sequentially. Although orientation estimates were more precise at the saccade target location, this benefit occurred only for the most recent of the two stimuli. Additionally, the more recent stimulus biased perceptual orientation estimates of the previous stimulus at the saccade target location. Analysis with a “variable precision” model^26,27^ revealed that incorporating both perceptual and choice biases was essential to fully account for the behavioral effects of presaccadic attention.

Our study uncovers a novel “presaccadic recency bias” that renders visual change detection challenging at the saccade target location. In conjunction with Bayesian modeling of behavior, these results identify the distinct contributions of perceptual and decisional processes to these behavioral effects of presaccadic attention. The finding – that the most recent information at the saccade target location is retained with high fidelity, and preceding stimulus percepts are biased towards the most recent – has critical implications for real-world scenarios (e.g. driving, sports) in which frequent gaze shifts occur in dynamically changing environments.

## Results

### Presaccadic attention enhances spatial choice bias but not sensitivity

To explore the effects of presaccadic attention on visual change detection, we developed a dual task paradigm that enabled quantifying change detection sensitivity and criterion at multiple locations on the screen as participants prepared and executed saccades. Briefly, on each trial, participants (n=10) were presented with 4 oriented Gabor patches, one in each visual quadrant, and cued to make a saccade to one of the four stimuli (25% probability across locations) as quickly and accurately as possible (Fig. 1A–B). A short, variable interval after the cue – but before the saccade was initiated – one (or none) of the Gabor stimuli underwent a change in orientation (Fig. 1A, top). Participants detected and localized the change with a 5-alternative response (Fig. 1A). Sensitivity and criterion were quantified from behavioral responses (Fig. 1D, left) with a well-validated signal detection model^24,25^ (Methods; Fig. 1D, right). The location cued for saccades was sampled independently from and was, therefore, not predictive of the location of change. This design enabled us to quantify the effects of presaccadic attention, without the confounding effects of voluntary (endogenous) spatial attention, on psychophysical parameters (sensitivity and criterion). Sensitivity and criterion were quantified at the saccade target location (ST or “toward”) and compared with their average value at the 3 other locations away from the saccade target (SA or “away”).

**Figure 1.**
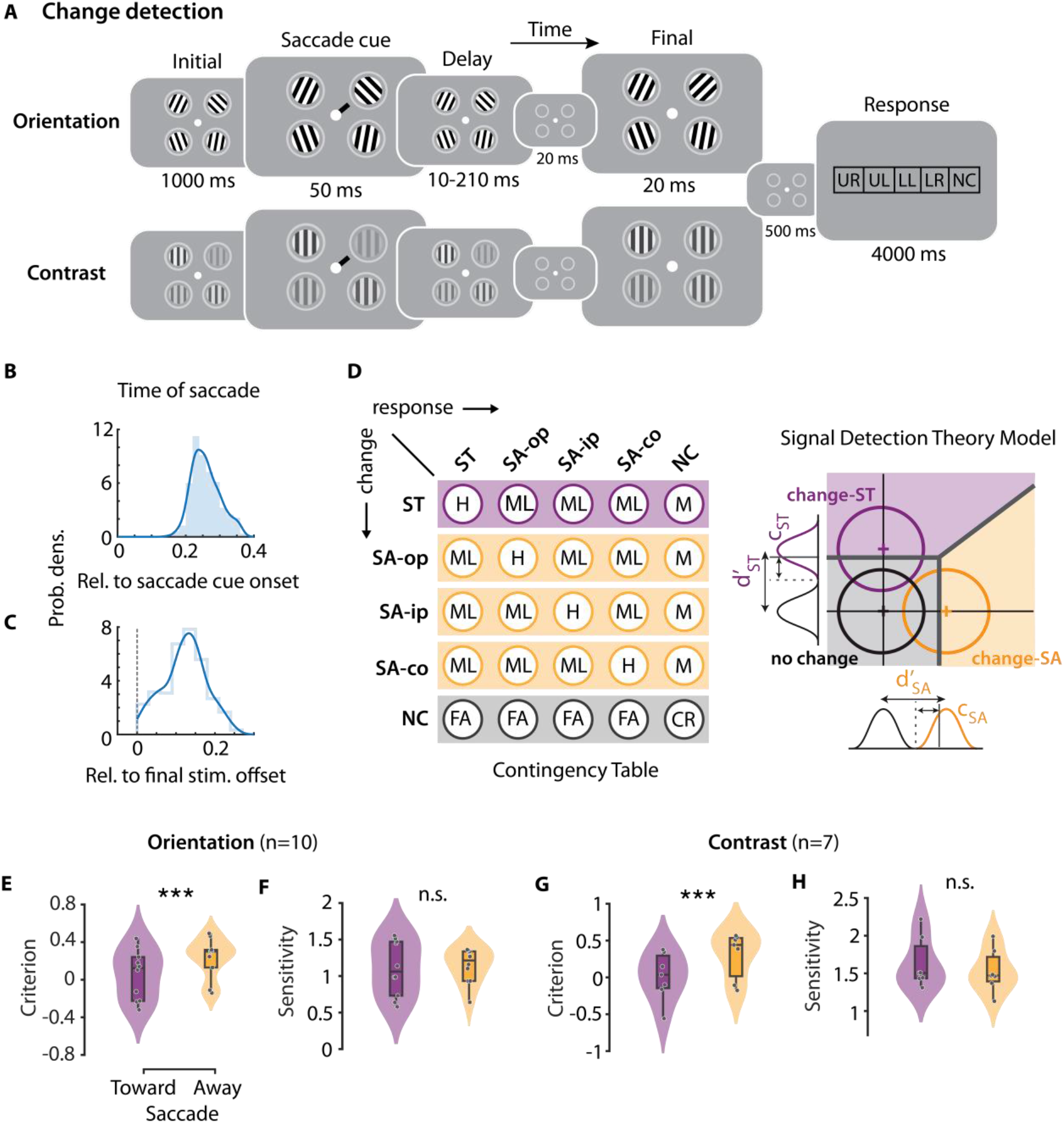
Presaccadic attention’s effects on visual change detection sensitivity and criterion. **A.** Dual task paradigm for quantifying presaccadic change detection performance in multi-alternative orientation change detection (top row) and contrast change detection (bottom row) tasks. Participants detected and localized changes in Gabor orientation (top) or contrast (bottom) while also planning a saccade to the cued location (see text for details). Rightmost panel: Response box configuration. UR: upper right, UL: upper left, LL: lower left, LR: lower right, NC: no change. **B.** Distribution (filled histogram) of the interval between saccade cue onset and saccade onset, across trials, for a representative (median) participant. Fit: kernel density estimate. **C.** Distribution (open histogram) of the interval between the final stimulus offset and saccade onset, for the same participant as in panel B. Other conventions are the same as in panel B. **D.** (Left) Stimulus-response contingency table for a multialternative change detection and localization (4-ADC) task. Rows: stimulus events, columns: response types. H: hits, M: misses, ML: mislocalizations, FA: false alarms, CR: correct rejections (see Methods for more details). ST: Saccade Toward; SA: Saccade Away; ip: ipsilateral, op: opposite, co: contralateral (all relative to the Saccade Toward location); NC: no change. (Right) Multidimensional signal detection model for estimating criterion and sensitivity at each location (see Methods for details). Change-ST: change at saccade target location (purple distribution and decision zone); Change-SA: change at saccade away location (orange distribution and decision zone); No change: no change at either location (black/gray distribution and decision zone). **E.** Criterion for orientation change detection reports at the saccade target (Toward) and non-saccade target (Away) locations (n=10 participants). Violin: rotated kernel density estimates; center line: median; box limits: upper and lower quartiles; whiskers: 1.5x interquartile range. All data points are shown. Purple: Saccade Toward (ST) location. Orange: Saccade Away (SA) location. **F.** Same as in panel B, but showing sensitivity for orientation change detection and localization. Other conventions are the same as in panel B. **G.** Same as in panel B, but showing criterion at the saccade Toward and Away locations (n=7 participants) for the contrast change detection task. **H.** Same as in panel F, but showing sensitivity for contrast change detection task. All panels. Asterisks: significance levels (* p<0.05, ***p <0.001). n.s.: not significant.

Given the extensive literature on presaccadic attention, we expected to find significant facilitation of behavioral metrics at the saccade target location. Surprisingly, change detection sensitivity was not higher at the saccade target location than at the other locations (Fig. 1F) (d’_Toward_=1.1±0.12, d’_Away_=1.15±0.08, p=0.529; random permutation test; psychophysical parameters averaged across the three away locations). On the other hand, change detection criteria were lowest – and spatial choice bias highest – at the saccade target location, relative to the other locations (Fig. 1E; c_Toward_=0.06±0.08, c_Away_=0.24±0.07, p<0.001). In other words, presaccadic attention enhanced spatial choice bias for change detection, but did not benefit detection sensitivity.

We performed additional control investigations to assess the validity and generality of these results. First, because the effects of presaccadic attention are transient^7,11^ we tested if sensitivity enhancements occurred in time windows proximal to the saccade. For this, we analyzed the dynamics of criterion and sensitivity changes in a restricted time window (−250 ms to 50 ms) locked to saccade onset. The variable interval between the saccade cue and the change event – as well as the natural variability in saccade onset times following cue presentation – enabled us to quantify sensitivity and criteria at different latencies of the change event relative to saccade onset. We observed strong, and significant, criterion modulations in multiple time windows (up to 200 ms) preceding saccade onset (SI Fig. 1A). On the other hand, sensitivity was not different between the saccade target location and the other locations in any time window tested (p>0.05 for all windows tested) (SI Fig. 1B).

Second, we tested if there was any difference in the effects on sensitivity or criterion at the non-saccade target locations ipsilateral (same hemifield) and contralateral (opposite hemifield) to the saccade target. Detection sensitivities were not significantly different between the ipsilateral and contralateral hemifield locations (d’_Away-Ipsi_=1.16±0.08, d’_Away-Contra_=1.14±0.10, p>0.99, random permutation test), and neither was significantly different from that at the saccade target location (d’_Toward_ vs. d’_Away-Ipsi_, p=>0.99; d’_Toward_ vs. d’_Away-Contra_, p=>0.99). Criterion at the contralateral locations was marginally higher than that at the ipsilateral location (c_Away-Ipsi_=0.16±0.08; c_Away-Contra_=0.28±0.06; p=0.004), yet each was significantly higher than that at the saccade target location (c_Toward_ vs. c _Away-Ipsi_, p=0.001; c_Toward_ vs. c _Away-Contra_, p<0.001).

Third, because in the main experiment we had tested a single orientation change angle – staircased independently for each participant (Methods) – we repeated this experiment with n=5 participants, comprising a subset of main cohort, by measuring the entire psychophysical function across multiple orientation change angles (Methods). Again, we observed that whereas detection criteria were reliably lowest at the saccade target location (p=0.002) (SI Fig. S1C), detection sensitivity was not significantly different between the saccade target location and the other locations (p=0.167) (SI Fig. S1D).

Fourth, we evaluated the generality of these results in another dual task paradigm involving contrast step detection performed by 7 participants, including 4 from the earlier cohort. The task design was identical to that of the main experiment, except that participants had to detect and localize a contrast increment – rather than an orientation change – that occurred in one of the four Gabor patches (Fig. 1A, bottom, Methods). Consistent with the main experiment findings, detection criterion was lowest at the saccade target location (Fig. 1G) (c_Toward_=0.01±0.12, c_Away_=0.31±0.12, p<0.001), but detection sensitivity was not significantly different across locations (Fig. 1H) (d’_Toward_=1.63±0.13, d’_Away_=1.53±0.11, p=0.307).

Finally, we quantified the strength of the observed effects across the change detection tasks, using the Bayes factor (BF; Methods). We estimated a BF of 10.18 for criterion effects and a BF of 0.32 for sensitivity effects (data pooled across orientation change and contrast change detection tasks). To confirm the robustness of these findings we performed Bayesian sequential analysis to check how evidence for the null versus the alternative hypotheses changed as data from successive participants were acquired (SI Figure S2A-B; see also Methods). These results provide strong evidence for a lower criterion at the saccade target location, and substantial evidence for no difference in sensitivity across locations.

To summarize, presaccadic attention produced no spatially specific improvement in change detection sensitivity. Rather, change detection criteria were lowest, and spatial choice bias highest, at the saccade target, presaccadically. In other words, presaccadic attention biased detection choices selectively at the saccade target location but did not improve the ability to detect changes at this location.

### A presaccadic recency bias renders change detection challenging

To further investigate the mechanisms underlying the surprising lack of presaccadic change detection benefits, we conducted another dual task experiment. Participants (n=10) estimated the orientation of one of four Gabor stimuli, presented presaccadically. The task structure was identical in nearly all respects to the saccade/orientation change detection dual task, except that participants reported their estimate of the orientation of the Gabor that was probed *post hoc*, by rotating a response bar (Fig. 2A) (Methods). Stimuli in the initial set or the final set were probed exclusively, in distinct trial blocks of trials (“double set” trials, Fig. 2A, middle). In a subset of randomly interleaved trials (40%), we replaced the final Gabor stimulus set either with a blank (n=5 participants) or with spatially filtered noise masks (n=5 participants). In the former trials only one set of stimuli was presented (“single set” trials; Fig. 2A, uppermost), whereas in the latter trials, the initial Gabor stimulus set was followed by a blank and noise masks (“noise mask” trials; Fig. 2A, lowermost).

**Figure 2.**
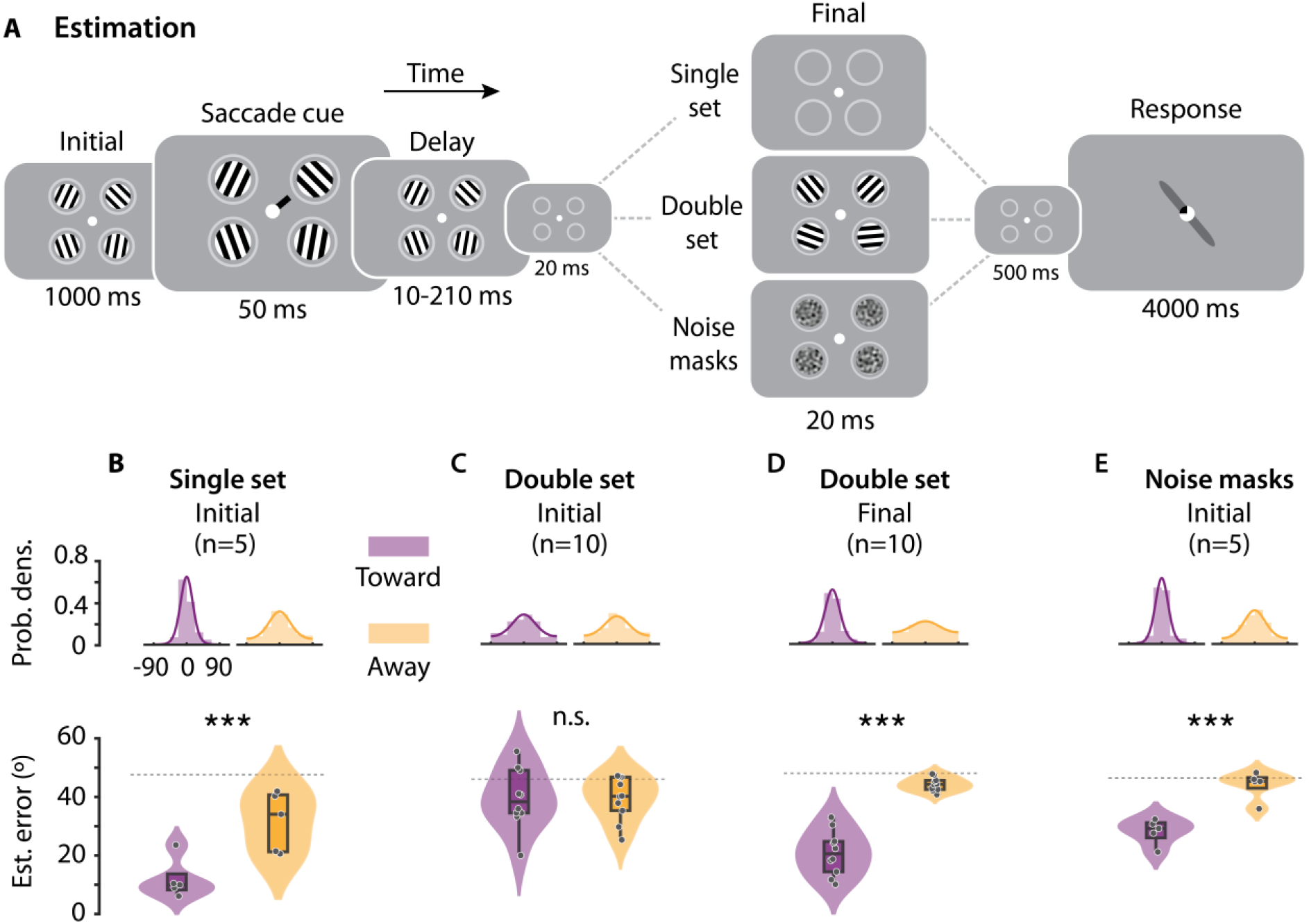
Presaccadic attention’s effects on precision of orientation estimates. **A.** Dual task paradigm for presaccadic orientation estimation, with three different trial types: (uppermost) single set trials, (middle) double set trials and (lowermost) noise mask trials Participants estimated the orientation of either the initial or the final set of Gabor stimuli in distinct blocks of trials, while also planning a saccade toward the cued location (see text for details). **B.** (Top) Distribution of estimation errors at the saccade Toward (left, purple) and Away (right, orange) locations for single set trials. Lines: von Mises fits. (Bottom) Mean absolute error (MAE) of orientation estimates at the saccade Toward (left) and Away (right) locations, for the initial stimulus of single set trials (n=5 participants). **C.** Same as in panel B, but showing error distributions and MAE for the initial stimulus of double set trials. **D.** Same as in panel B, but showing error distributions and MAE for the final stimulus of double set trials. **E.** Same as in panel B, but showing error distributions and MAE for the initial stimulus of noise mask trials. (B-E). Other conventions are the same as in Figure 1E.

Single set trials revealed systematic benefits of presaccadic attention: Orientation estimation accuracy was significantly higher, and absolute error of orientation estimates (“estimation error”) nearly 2x lower, at the saccade target location than at the other, non-saccade target, locations (Fig. 2B) (mean±s.e.; Err_Toward_=14.4°±2.6°, Err_Away_=31.5°±3.9°, p<0.001, permutation test; BF=22.51); similar results were obtained with quantifying the precision (Methods) of orientation estimates (Fig. 2B, inset; Precision_Toward_=1.30±0.39, Precision_Away_=0.66±0.08; p<0.001, random permutation test). In other words, when only one stimulus was presented at the saccade target location, presaccadic attention improved the precision of orientation estimates, in line with results reported in multiple previous studies.

Yet, in double set trials – when the initial set of stimuli was followed by a second (final) set – this presaccadic benefit vanished entirely for the initial Gabor stimulus. Estimation error for the initial Gabor stimulus was not different between the saccade target location and the other locations (Fig. 2C) (Err_Toward_=39.5°±2.8°, Err_Away_=39.5°±2.0°, p=0.526, random permutation test; BF=0.3). Remarkably, a significant presaccadic benefit occurred, now, for the most recent Gabor stimulus: estimation error for the final Gabor stimulus was lowest at the saccade target location, as compared to the other locations (Fig. 2D) (Err_Toward_=21.7°±2.1°, Err_Away_=41.9°±0.6°, p<0.001; BF>10^3^). In other words, when two sets of stimuli were presented in sequence, presaccadic benefits occurred only for the last of the two stimuli.

We tested whether the loss of precision could be accounted for by visual “masking” of the initial stimulus by the final stimulus. For this, we presented spatially filtered noise masks, rather than Gabor stimuli, in the final set (Fig. 2A, lowermost, Methods). Interestingly, in this case, presaccadic benefits persisted at the saccade target location (Err_Toward_=29.3°±1.7°, Err_Away_=43.0°±1.8°, p<0.001, permutation test; BF=32.98) (Fig. 2E). In other words, sensory masking effects could not explain the loss in precision for the initial stimulus in the double set trials. Rather, the final stimulus must be task relevant – in this case, an oriented Gabor stimulus – to abolish presaccadic benefits for the initial stimulus.

To investigate these temporal order effects further we tested whether, and how, the percept of the initial stimulus at the saccade target location was influenced by the final stimulus. Indeed, orientation estimates of the initial stimulus were strongly and systematically biased (Methods; Fig. 3A) toward the orientation of the final stimulus, even though the initial and final Gabor stimuli were uniformly and independently sampled (Fig. 3B–C) (Bias=25.7±2.6, p=0.002, Wilcoxon signed rank test; BF>10^3^). In contrast, the orientation estimates of the final stimulus were not biased by the orientation of the initial stimulus (Fig. 3D–E) (Bias=-2.7±1.3, p=0.193; BF=1.37). Interestingly, at the non-saccade target locations we observed a converse trend: orientation reports of the final stimulus were marginally biased toward the initial stimulus orientation (Bias=7.4±2.3, p=0.014; BF=5.68), but not vice versa (Bias=4.0±2.6, p=0.131; BF=0.76). A detailed Bayesian sequential analysis robustness check is provided in SI Figure S2C-F.

**Figure 3.**
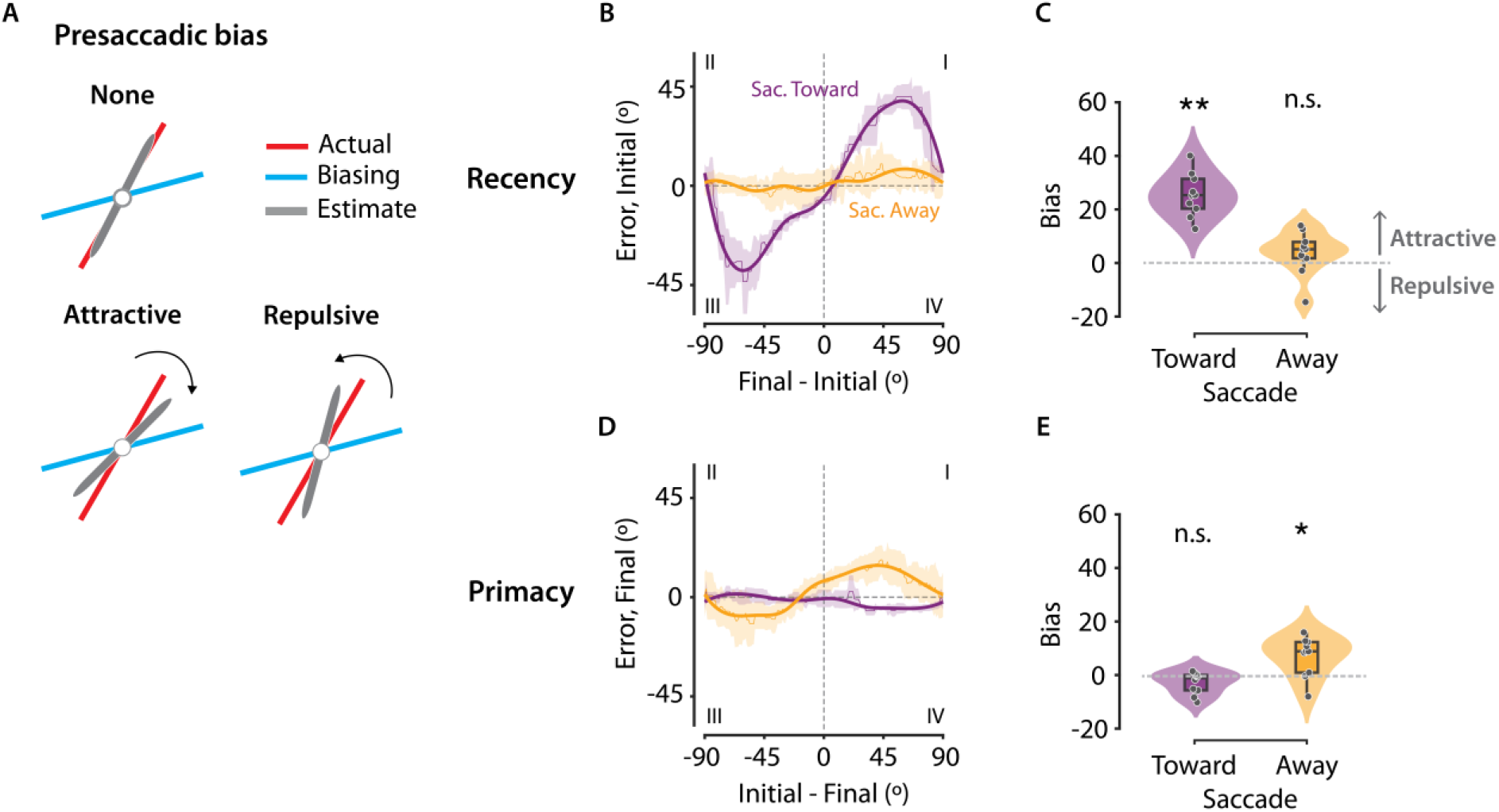
Evidence for a presaccadic recency bias in orientation estimation. **A.** Schematic showing an attractive bias (bottom, left), a repulsive bias (bottom, right) and no bias (top). In the case of an attractive bias, orientation estimates (gray) of the actual stimulus (red) are biased toward the orientation of the biasing stimulus (blue). In the case of the repulsive bias, orientation estimates are biased away from the biasing stimulus. **B-C.** Recency bias: the bias in the orientation estimate of the initial stimulus induced by the final stimulus. **B.** Estimation error for the initial stimulus (y-axis) plotted against the difference between orientations of the final and initial stimuli (x-axis), at the Saccade Toward (purple) and the Saccade Away (orange) locations. Dashed horizontal and vertical lines: zero estimation error and zero orientation difference, respectively. Thick, solid lines: Cubic spline fits. **C.** Recency bias at the Saccade Toward (left, purple) and Away (right, orange) locations in the double set trials, quantified using the signed area under the curve, from panel B (Methods). Positive and negative values denote an attractive or a repulsive bias, respectively. ** p<0.01. Other conventions are the same as in Figure 1E. **D–E.** Primacy bias: the bias in the orientation estimate of the final stimulus induced by the initial stimulus. **D.** Same as in panel B, but showing the estimation error for the final stimulus (y-axis) plotted against the difference between orientations of the initial and final stimuli (x-axis). Other conventions are the same as in panel B. **E.** Same as in panel C, but showing primacy bias. Other conventions are the same as in panel C.

We tested if these effects were not due to presaccadic temporal order reversal effects – an illusory reversal in sequence order of a sequence that occurs ~50 ms before a saccade^28,29^. We re-evaluated the presaccadic recency and primacy biases by excluding trials where the offset of the final set of stimuli was within 50ms of the saccade onset (SI Fig. S3A- B). The estimates of the initial stimulus continued to be biased towards the final stimulus at the saccade Toward location (Bias=24.6±2.7, p=0.002), confirming that the presaccadic recency bias was not due to an apparent confusion in the stimulus order.

To summarize, the precision of orientation estimates was highest at the saccade target location, but only for the more recent of the two stimuli presented at this location. Moreover, orientation estimates for the initial stimulus were biased toward the orientation of the recent stimulus. This presaccadic “recency” bias, therefore, rendered accurate change detection at the saccade target location potentially even more challenging.

### Both perceptual and choice biases contribute to presaccadic attention’s effects

Although the recency bias could explain the lack of presaccadic benefits on change detection sensitivity, it appears at odds with our finding regarding a higher spatial choice bias for reporting changes at the saccade target location. Because of the recency bias, the perceived difference in orientation between the initial and final Gabors would always be smaller in magnitude than the actual difference. This would yield fewer false alarms, a higher criterion, and, therefore, a lower bias at the saccade target location; whereas we observed the opposite trend – a lower criterion, and higher bias – at the saccade target location.

We sought to reconcile these apparently contradictory findings using a Bayesian ideal observer model: the variable-precision (VP) model^26,27^. The VP model has been successfully applied for modeling behavior in multialternative choice tasks^26,27^; in our study, it provided a unified framework for modeling behavior both in the orientation change detection task (Fig. 1) and the orientation estimation task (Fig. 2). Briefly, the model posits that neural resources, and, therefore, the precision of stimulus representation, are variable across items and trials. The model estimates two parameters, 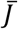 and *τ* (scale), from which the mean and the variance of the stimulus precision can be computed (as 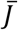 and 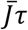, respectively, Fig. 4A) (Methods). Specifically, we tested whether one or both types of presaccadic bias – the perceptual (recency) bias, observed in the estimation task or the decisional (choice) bias, observed in the change detection task – would be relevant for explaining presaccadic attention’s effects.

**Figure 4.**
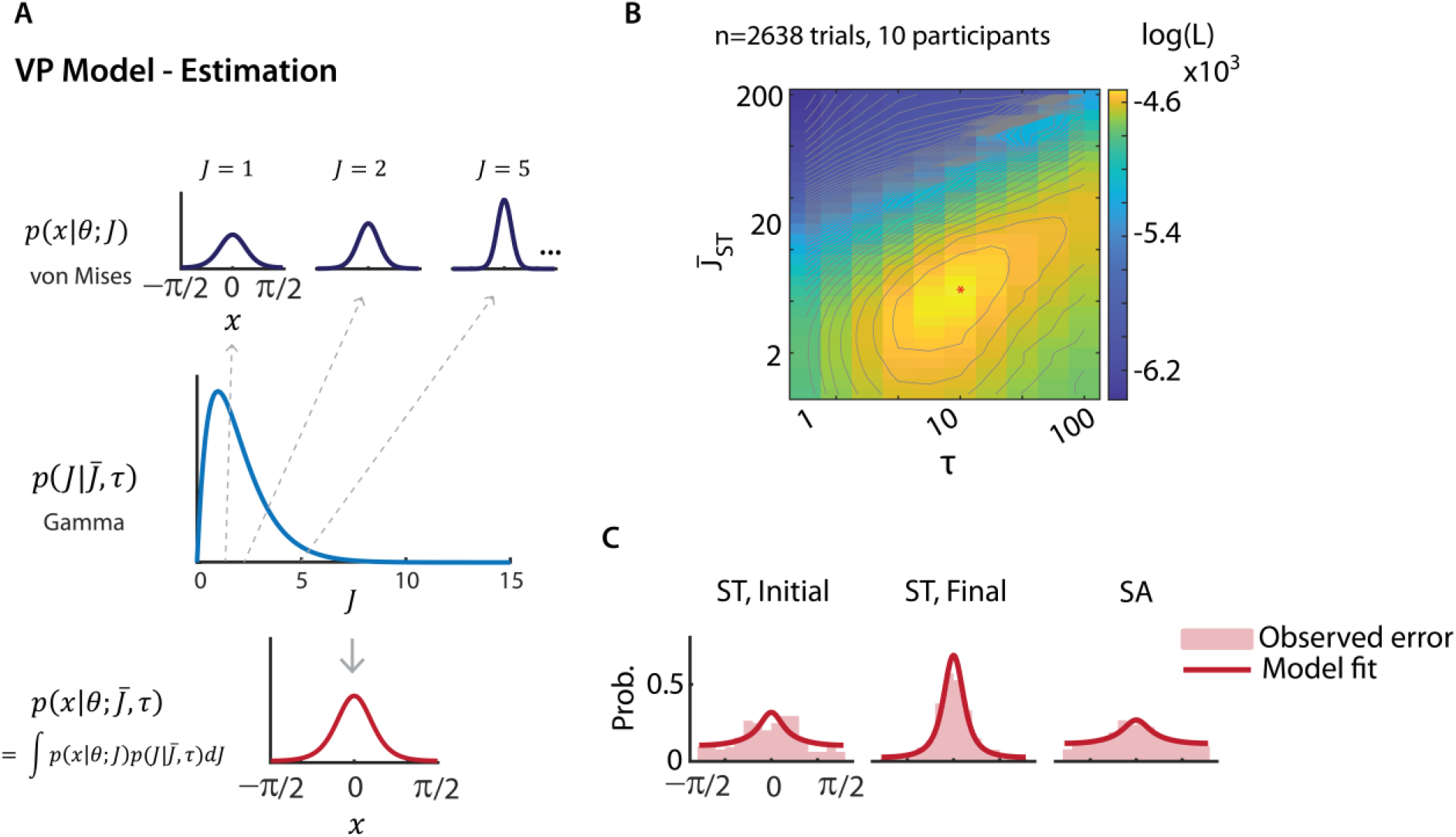
Variable precision model fits presaccadic attention’s effects on orientation estimation. **A.** Schematic of the variable precision model for orientation estimation. (Top row) On each trial, internal measurement, x, of a stimulus with orientation, θ, follows a von Mises distribution, with a concentration parameter κ determined by precision parameter J. (Middle row) Across trials the precision is gamma distributed, with a mean 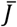 and scale parameter τ. (Bottom row) The distribution of the internal measurement x across trials is a mixture of von Mises distributions, marginalized across precision values. **B.** Log likelihood (log (L)) landscape as a function of precision for the final stimulus at the Saccade Toward location 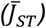 and scale, τ; although log likelihood is a function of all model parameters, these two parameters were chosen for illustrative purposes. Warmer colors indicate higher likelihoods. Red asterisk: Parameters corresponding to maximum log likelihood. Gray lines: contours of iso-log likelihood. **C.** Left to right: Fits of the variable precision model (solid red lines) to the observed responses (light red histograms) for the initial ST, final ST and SA (pooled) stimuli, respectively, obtained with maximum likelihood estimation, based on the parameters in panel B.

First, to limit the number of parameters required to fit the orientation change detection task, we identified *τ* (scale) *apriori* by fitting the VP model to response error distributions from the orientation estimation task. Based on our empirical findings (Fig. 2C–D), we employed 3 different mean precision parameters: two to model the different precisions for the initial and final stimuli at the saccade target location, and one to model the precision for both stimuli at the non-saccade target locations. In line with experimental observations, the VP model estimated a significantly higher mean precision for the final ST stimulus, as compared to both the initial ST stimulus as well as both set of SA stimuli (Fig. 4B) (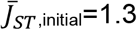, 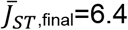, 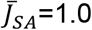; p<0.001, permutation test, data pooled across subjects). In addition, *τ* was estimated to be 13.0. Visualizing the fit of the VP model to the error distributions (Fig. 4C), revealed that the model successfully captured the effect of presaccadic attention on the precision of orientation estimates (p>0.1, Kuiper’s test).

Next, we fit the VP model to the stimulus-response contingency table in the multialternative change detection task (Fig. 5A) using a variant of the VP model^26^ – a change localization model – that, on each trial, reports the location with the highest posterior probability of change occurrence. To model change detection, we modified the decision rule by incorporating a “decision threshold” for the posterior probability, below which the model provides a no-change response (Methods). We tested three variants of this model: i) a “baseline” (standard VP) model that modeled unequal precisions at the ST and SA locations but imposed a common decision threshold of 1.0 across locations (Fig. 5A, black); ii) a “perceptual bias” model that extended the baseline model by additionally estimating a recency bias for the initial stimulus representation toward that of the final stimulus at the saccade target location (Fig. 5A, red); iii) a “both biases” model that extended the perceptual bias model by additionally modeling choice bias, viz., different decision thresholds at each location (Fig. 5A, blue). Models were formally compared with the corrected Akaike Information Criterion (AICc) and cross-validated data likelihoods.

**Figure 5.**
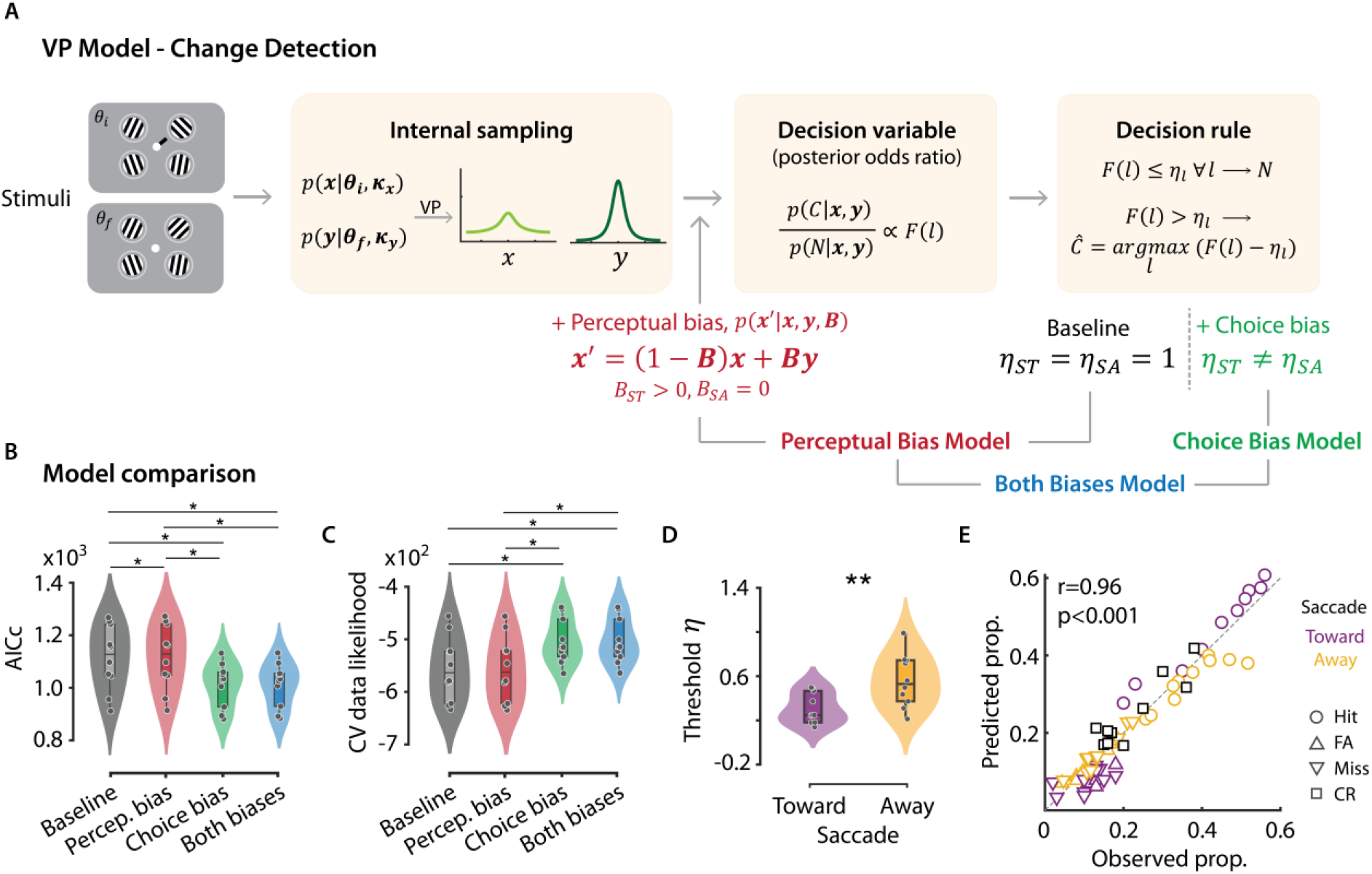
Perceptual and choice biases contribute to presaccadic attention’s effects on orientation change detection. **A**. Schematic of the variable precision model for a change detection and localization task. (Left box) Internal measurements of the initial (**θ_i_**) and final (**θ_f_**) Gabor stimuli are represented by latent vectors (**x** and **y**, respectively) that follow von Mises distributions, with concentration parameters, **κ**. The encoding precision (**J**, directly related to **κ**) is variable across items and trials. (Middle box) F(l), the decision variable at each location, l, is proportion to the posterior odds ratio of probability of change (C) at that location to the probability of no change (N). The Bayesian ideal observer localizes the change to the location at which the decision variable F(l) exceeds the decision threshold by the highest margin. If the decision variable does not exceed the decision threshold at any location, the observer reports “no change”. Perceptual bias model: The “baseline” VP model was modified to incorporate a “perceptual bias”, i.e., an attractive recency bias parameter, B, at the Saccade Toward location (red text). Choice bias model: The “baseline” VP model was modified to incorporate unequal decision threshold parameters, η_l_, at the Saccade Toward and Saccade Away locations (green text). Both biases model: Both perceptual bias and unequal decision threshold parameters were incorporated into the VP model (blue text). **B**. Model comparison with the Akaike Information Criterion (AICc), for the four VP models. Gray/black: baseline model, red: perceptual bias model, green: choice bias model, blue: both biases model. Box-and-whisker plot conventions are the same as in Figure 1E. **C**. Same as in panel B, but showing the cross-validated data likelihoods for the four models. Other conventions are the same as in panel B. **D**. Same as in Figure 1E, but showing the decision threshold, η, estimated for the Saccade Toward and Saccade Away locations with the “both biases” model. Other conventions are the same as in Figure 1E. **E**. Response proportions predicted with the both biases model (y-axis) versus observed (true) response proportions (x-axis) in the change detection task. Circles: hits, upward triangles: false-alarms (FA), downward triangles: misses, squares: correct rejections (CR). Other conventions are the same as in panel D.

In line with expectations the perceptual bias model did not improve baseline model fits (Fig. 5B; median AICc_baseline_=1128.5±37.6; AICc_perceptualBias_=1130.5±37.7, p=0.012). In fact, AICc values were marginally worse for the perceptual bias model: simply adding a recency bias at the ST location would yield lower (not higher) false-alarms at this location, as noted above. In addition, evaluating the goodness-of-fit based on a randomization test (Methods) revealed successful fits for none of the 10 participants with either the baseline model or with the perceptual bias model. In stark contrast, upon incorporating differential decision thresholds at each location AICc values improved significantly (Fig. 5B, right; AICc_bothBiases_=1030.9±26.0, p=0.012). Moreover, the “both biases” model estimated a systematically lower decision threshold for the ST, as compared to the SA, location (Fig. 5D, *η*_ST_=0.31±0.04, *η*_SA_=0.56±0.06, p=0.002).

To confirm that the more complex model was not over-fitting the data we evaluated cross-validated data likelihoods (Methods). Again, the cross-validated likelihoods were highest for the model that incorporated both biases as compared to either the perceptual bias or baseline models (cvL_baseline_—562.6±8.8, cvL_perceptualBias_—561.4±18.9, cvL_bothBiases_—513.3±13.3) (Fig. 5C). The both biases model successfully fit all 10/10 participants’ data (median gof p-value: 0.87): the predicted responses closely matched the observed responses in the model that incorporated both biases (r_bothBiases_=0.96, p<0.001) (Fig. 5E) as compared to the other two models (r_baseline_=0.45, r_perceptualBias_=0.44, SI Fig. S4A-B).

A model that incorporated only choice bias – and no perceptual bias – was only marginally worse than the both biases model (AICc_choiceBias_=1034.1±26.3; cvL_choiceBias_=- 514.4±13.3; median gof p-value: 0.83). Yet, as compared to the “both biases” model, median AICc for the “choice bias” model was higher by 3.45 points, cross-validated likelihoods were worse for 7/10 participants, and goodness-of-fit p-values were poorer for 7/10 participants, suggesting that modeling perceptual bias (along with choice bias) contributed to improving model fit to behavior.

In sum, Bayesian analysis with the VP model revealed the role of perceptual and choice biases in mediating the behavioral effects of presaccadic attention. Although the perceptual recency bias was useful for explaining presaccadic attention’s effects on sensitivity, choice bias was arguably the single most important factor for presaccadic effects on criteria, and significantly improved model fits to behavior. Taken together, our results pinpoint dissociable mechanisms by which perceptual and decisional processes mediate presaccadic attention’s effects on behavior. Moreover, they provide a parsimonious account for why presaccadic benefits – reported widely in visual discrimination tasks – are surprisingly absent during visual change detection.

## Discussion

Planning to move the eyes changes the way we perceive our visual world^2,3,12,28,30–34^. Many previous studies have shown that presaccadic attention enhances visual discrimination accuracy, immediately before the eyes move; several more studies have proposed specific behavioral and neural mechanisms underlying these benefits^6,13–15,31–33^. In a surprising counterpoint to these earlier studies, we show that presaccadic attention does not benefit visual change detection: while, presaccadic attention biased change detection choices toward the saccade target location it produced no effect on detection sensitivity.

We explored this surprising result further with a presaccadic orientation estimation paradigm, that involved estimating the orientation of one of two Gabor patches, presented in quick succession. While presaccadic attention did enhance the precision of orientation estimation, this benefit was observed only for the most recent stimulus at the saccade target location. Moreover, the results provide evidence for a novel “presaccadic recency bias” – a bias with reporting the orientation of the initial stimulus induced by the final stimulus at the saccade target location.

The presaccadic recency bias could readily explain the absence of change detection sensitivity benefits at the saccade target location: because the perceived orientation of initial stimulus was biased towards the final stimulus, accurate computation of the orientation change was likely compromised at this location. Yet, the enhanced precision for the final stimulus possibly compensated for this detriment in the change signal computation. The net effect was, therefore, a lack of a robust enhancement in visual change detection sensitivity at the saccade target location, relative to the other locations.

An alternative explanation for the presaccadic recency bias is the illusory temporal order reversal that occurs for sequences presented presaccadically^28,29^. We tested, and discounted, this explanation. For one, this reversal effect has been described only for highly transient stimulus sequences (~10 ms flashes separated by 100 ms), and for reflexive saccades. Given that, in our task, the first stimulus set was displayed on screen for >1000 ms, and the participants made volitional saccades, such a temporal order reversal is unlikely to have occurred. Second, the effect of temporal reversal has been described perisaccadically, in a time window spanning ± 50ms locked to saccade onset. In our experiments, even after excluding trials within this range, we continued to observe the presaccadic recency bias (SI Fig. S3A). Finally, while reports for the initial Gabor stimulus orientation were indeed closer to the final stimulus orientation than their own, the converse was not true; these results demonstrate clearly that participants were not mistakenly interchanging the identities of the initial and final Gabors.

At first glance, the recency bias appears to be at odds with our finding regarding criteria at the saccade target location: Because of the recency bias, the perceived difference in orientation between the initial and final Gabors will always be smaller in magnitude than the actual difference. This should yield fewer false alarms, and a higher criterion, whereas we observed the converse – a lower criterion – at the saccade target location. These observations could be reconciled if we posit that in addition to the perceptual recency bias, presaccadic attention also produced a decisional (choice) bias – that reduced the decision criterion – at the saccade target location. Such a choice bias must overwhelm the effects of the perceptual bias, thereby yielding the highest proportion of false alarms at the saccade target location.

We tested this hypothesis using a Bayesian variable precision model. The variable precision model simulates trial-wise variations in internal estimates of orientation, based on which a Bayesian ideal observer would perform change detection and localization. In our case, the model also offered an elegant way of modeling both change detection and estimation tasks within a single framework. The VP model variant that best fit behavior was one that not only accounted for differences in precision across locations but also incorporated both a choice bias (lower detection threshold) and an attractive presaccadic recency bias at the saccade target location. In addition, given the potential for overfitting with the multi-parameter VP model, we estimated cross-validated data likelihoods using behavioral predictions of left-out sessions, which confirmed the robustness of these findings.

While previous studies have used elegant task designs to study behavioral and neural mechanisms of presaccadic attention on visual sensitivity^3,6,7,9,12,20,23^, none, to our knowledge has systematically explored the effects on spatial choice bias. Neural evidence indicates that a presaccadic shift and scaling of visual receptive fields^30,32,35,36^ at the saccade target may underlie the enhanced visual discrimination sensitivity prior to saccades. Furthermore, it has been shown that the presaccadic response of the visual cortex preferentially encodes the stimulus features at the saccade target^31^. Our results are consistent with such a mechanism by which the most recent stimulus is selected for processing with the highest fidelity. Moreover, in many studies^7,9,20^, a single discrimination target was displayed among distractors without an explicit location probe; such a design precludes inferring the specific stimulus that influences the participant’s report. Moreover, biases reported in visual discrimination tasks reflect biases for different stimulus features (e.g. clockwise versus counter-clockwise orientations). By contrast, the change detection and localization response in our current design – along with analysis with a signal detection theory model^24^ – enabled precisely decoupling and quantifying location-specific effects of presaccadic attention on sensitivity and bias. The presaccadic gain modulation in neural responses in the visual cortex^30,32,33^ is consistent with an increase in choice bias for selecting the saccade target for response; similar bias (criterion)- related gain modulations have been reported previously in visual cortex^37^.

Our results add to a growing body of literature that investigates shared and distinct mechanisms mediating covert endogenous attention and presaccadic attention. A few previous studies^6,17^ employed a saccade cue that was 100% predictive of the location of the upcoming discriminandum; such a design may conflate the effects of presaccadic attention with those of covert endogenous attention. But in our study the saccade cue was entirely uninformative about the upcoming location of change; this design enabled us to distinguish the effects of presaccadic attention from those of endogenous attention. Whereas endogenous attention has been reported to enhance both change detection bias and sensitivity at the location cued for attention^38–41^, we show that presaccadic attention enhances change detection bias, but not sensitivity, at the saccade target location. Given that we employed a task design with stimulus configuration and timings closely similar to previous endogenous attention studies, we propose that these differences in behavioral effects between these types of attention arise due to the unique nature of the presaccadic recency bias. While the recency bias occurred in this presaccadic attention task – and impacted change detection sensitivity – no such bias has been reported in endogenous attention tasks. Identifying neural substrates underlying sensitivity and criterion modulation by endogenous attention, as well as those underlying the recency bias for presaccadic attention may help identify shared and distinct mechanisms underlying these two types of attention.

Our experiments can be readily extended in many directions. While we report an all-or-none presaccadic benefit that occurs entirely for the last of two stimuli, and not at all for the first, it would be interesting to know if the effect occurs in an all-or-none or graded manner when a sequence of multiple stimuli is presented presaccadically. Furthermore, investigating presaccadic recency bias in a naturalistic setting (e.g. with VR displays) may inform our understanding of how a stable percept of the world emerges despite frequent eye movements in free-viewing, real-life conditions.

## Acknowledgements

This research was supported by a Wellcome Trust-Department of Biotechnology India Alliance Intermediate fellowship, DST Swarna-Jayanti fellowship, a Pratiksha Trust Intramural grant, an India-Trento Programme for Advanced Research (ITPAR) grant and a Department of Biotechnology-Indian Institute of Science Partnership Program grant (all to DS).

## Materials and Methods

### Participants

A total of 21 unique participants (9 females, age range: 19-36 years, median age: 24 years, all right-handed), with normal or corrected-to-normal vision, and no known history of neurological disorders participated in the experiments; one of the authors (PG) also participated. 10 participants (6 females, age range: 19-27 years) performed the orientation change detection task. 7 participants (2 female, age range: 19-27 years) performed the contrast increment detection task. 10 participants (4 females, age range: 20-36 years) performed an orientation estimation task. 4 participants performed both the orientation change detection and contrast increment detection tasks. 1 participant (2 participants) performed both the orientation change detection (contrast increment detection) and orientation estimation tasks. Informed written consent was obtained from all participants, and experimental protocols were approved by the Institute Human Ethics Committee, Indian Institute of Science, Bangalore.

### Experimental design

#### Behavioral Tasks

##### Setup

Participants performed the tasks in a dimly-illuminated room, with their head stabilized on a chin and forehead rest, seated 60 cm from a contrast-calibrated stimulus display (BenQ XL2411Z, 100 Hz refresh rate). The task was designed with custom scripts using Psychtoolbox (version 3.0.15) implemented on MATLAB 2015b. Gaze position was tracked with an infrared eyetracker (SMI iViewX HiSpeed, SensoMotoric Instruments) at a 500 Hz sampling rate; analyses of gaze position were performed both online and offline (see next).

##### Orientation change detection and localization task

Participants performed a dual task, comprising detecting and localizing orientation changes at one of four peripheral Gabor patch stimuli (4-ADC task^24^), while also planning and executing saccades to one of the four stimulus locations. A trial began with fixation on a central dot (positive contrast, 0.3° diameter) concurrently with the presentation of four circular placeholders (positive contrast circles, 5.3° diameter), one in each visual quadrant at an eccentricity of 5.8° along the diagonal. Centered within each placeholder a Gabor patch (30% peak contrast, Gaussian s.d.: 0.8°, spatial frequency: ~1.9 cycles/°, aperture diameter: 4.7°) was presented. Gabor orientations were drawn from circular uniform distributions, independently of each other. Following 1000 ms of this initial stimulus set, a saccade cue (line segment, negative contrast, radial extent: 0.3°) adjoining the central dot appeared for 50 ms, pointing towards one of the quadrants. Participants were instructed to make a saccade towards the center of the stimulus in the cued quadrant (“saccade target”) as accurately and as soon as possible following saccade cue onset. Following saccade cue onset, the initial stimulus set remained on the screen for a variable, additional interval drawn from a geometric distribution (10-210 ms, in steps of 20 ms). Following this, the stimuli disappeared for 20 ms, and reappeared for 20 ms with either one or none of the Gabor stimuli having changed in orientation. Upon successful completion of the saccade (gaze position crossing the appropriate placeholder boundary) a set of five response boxes appeared on the screen in a linear row (Fig. 1A). Participants indicated the location at which they perceived a change in orientation or “no change” by clicking the appropriate response box with an optical mouse (4000 ms response window). This linear layout of response options was chosen so as to avoid systematic spatial correspondences between the location of the response buttons and the saccade target location, either in terms of left versus right, or upper versus lower, hemifield locations (Fig. 1A); the goal was to avoid response biases due to eye-hand coordination – which may occur if the location selected for oculomotor response were aligned with the subsequent manual response. The response boxes also appeared if no successful saccade was recorded within 400 ms of saccade cue onset; such trials (mean ± s.d. across participants: 11.9% ± 8.9%) were discarded before subsequent analyses. The saccade cue was not informative about the location of change: the change was equally likely to occur at any of the four locations (20% trials), or not at all (20% trials). Participants were informed about these task design elements, *a priori.*

The experiment was conducted over two days. On the first day participants were trained on the 4-ADC task with a 90° change angle, without a saccade cue, but with feedback on their accuracy on each trial. Following this, the change angle was staircased for each participant to ~55% accuracy, also with the no-saccade 4-ADC task (mean ± SD: 17.70 ± 4.08°). Following the staircasing, participants were trained on the dual task paradigm in which they performed the change detection along with saccade execution, and were provided with feedback for both task objectives. This training was typically performed for 75 to 150 trials, with the staircased change angle.

On Day 2, in a preliminary training session, participants performed 1-4 blocks of 75 trials each, after which testing commenced. The testing phase comprised 8 blocks of 75 trials without feedback as part of the main experiment. In a subset of 5 participants, we measured the psychometric function with four change angles (10°, 25°, 40°, 55°); these sessions, comprising 800 trials per participant, were acquired on a separate experimental day. Data from the training blocks were not included in the subsequent analyses.

##### Contrast increment detection and localization task

Participants were tested on a dual task, as before, except that they detected and localized a contrast increment at one of four Gabor stimuli while also planning and executing saccades to one of the four stimulus locations. The task design was similar to the orientation change detection task, except for a few differences in stimulus configuration. First, all of the Gabor stimuli were vertically oriented, and their contrasts were drawn, randomly and uniformly, from four non-overlapping contrast bins ([10-20%], [30-40%], [50-60%], [70-80%]). Second, when the stimuli reappeared after the first blank (Fig. 1A, bottom), the contrast of one or none of the stimuli was incremented by 20%. Participants were instructed to indicate the location of perceived increase in contrast, or no change, as before, by clicking on one of five response boxes. Finally, staircasing was performed by varying the duration of the final set to achieve ~55% accuracy (30-60 ms across participants).

##### Orientation estimation task

Participants performed a dual task, as before, except that they estimated the orientation of one of four Gabor stimuli while also planning and executing saccades to one of the four stimulus locations. Stimulus sequence and timings were nearly identical with the orientation change localization task except that the initial and final Gabor orientations at each location were drawn independently from a circular uniform distribution (Fig. 2A, middle). In addition, the initial stimulus set duration was drawn from a geometric distribution (10-210 ms, steps of 100 ms). In distinct blocks of trials, participants estimated the orientation of the probed Gabor either from the initial or from the final set of stimuli; written instructions regarding the set being probed (“initial set probed” or “final set probed”) were provided onscreen at the beginning of each block. At the end of the trial, a central response probe (positive contrast central dot, 0.5° diameter, with negative contrast quadrant) indicated the location of the Gabor stimulus for response. Upon moving the mouse cursor, a randomly oriented grey bar (2.9° length, 0.3° thickness) appeared behind the central probe. Participants were instructed to rotate the response bar until the bar’s orientation matched that of the probed Gabor, and clicked the left mouse button to complete their response. Participants were afforded 1500 ms to initiate and 2500 ms to complete their response. As with the previous tasks, the saccade cue was uninformative regarding the location of the probed Gabor grating.

In 60% of trials, Gabor stimuli were presented both as the initial and final set of stimuli “double set” trials). In one group of (n=5) participants, in 40% of trials interleaved with the double set trials, only the initial set of Gabor stimuli were presented, and the second set of Gabor stimuli were replaced with a set of blanks (“single set” trials). In the remaining (n=5) participants, in 40% of trials, interleaved with the double set trials, the second set of Gabor stimuli were replaced with filtered noise masks (“noise mask” trials). Noise masks were generated by bandpass filtering Gaussian noise (mean background contrast, SD: 16.7% contrast) in a band from 0.5x to 2x the spatial frequency of the Gabor stimuli^6^. Because only one set of Gabor stimuli were presented in both single set and noise mask trials, for these trials participants were instructed to report the orientation of whatever Gabor stimulus they perceived at the probed location, regardless of block type (initial set probed or final set probed).

#### Eyetracking and trial exclusion

Gaze position was calibrated with a 9-point calibration (iViewX) at the beginning of the experiment, and periodically throughout the experiment. Saccade onsets were detected offline with the BeGaze software based on the following criteria: gaze deviation from fixation >2°, velocity >50 °/s, duration >8 ms, and confirmed with manual inspection. For subsequent analyses, the following stringent criteria were used for trial inclusion based on gaze dynamics: i) the saccade must be made to the correct quadrant, as indicated by the saccade cue; ii) gaze position must cross the placeholder boundary within 400 ms of saccade cue onset; iii) stable fixation (<1° change in gaze position) from saccade cue onset, until saccade onset. In addition, we excluded those trials in which the saccade onset occurred before the second set of stimuli had disappeared. Based upon these criteria the trial inclusion rates for the different tasks were as follows (mean ± SD, across participants): i) orientation change detection and localization: 78.7% ± 6.4%; ii) contrast increment detection and localization: 73.7% ± 6.7%; iii) orientation estimation: 64.8% ± 7.2%.

### Data Analyses

#### Change detection and localization tasks

For the change detection and localization tasks, we estimated psychometric and psychophysical parameters at the ST and SA locations. The same analysis approach was used for both the orientation change detection and the contrast increment detection tasks.

##### Constructing contingency tables

Prior to signal detection theory analysis, we constructed 5×5 stimulus-response contingency tables for each participant using all trials from each experimental session (Fig. 1D, left). Rows of this table represent the five potential stimulus events (change at each of the 4 stimulus locations, or no change), whereas columns represent the five possible responses (report of change detected at each of the 4 locations, or no change). For these analyses, trials were sorted so that the first 4 rows of the table represented the location of the change relative to the saccade cue: cued (location towards which the saccade cue pointed), opposite (location diametrically opposite to the cued location), ipsilateral (location in the same hemifield as the cued location), and contralateral (location in the opposite hemifield as the cued location). We refer to the cued location as the “saccade toward” or “saccade target” (ST) location, and the opposite, ipsilateral and contralateral locations collectively as the “saccade away” (SA) locations. Similarly, the first 4 columns were re-sorted so that they represented the participant’s response location relative to the saccade cue. Then contingency table encodes response proportions for multiple different response types (Fig. 1D, left). The values along the diagonal encode correct responses: hits (H), when the participant reported the location of change accurately and correct rejections (CR) when the participant correctly reported no-change. The off-diagonal elements encode three kinds of incorrect responses: misses (M) when the participant reported no-change even when a change occurred (last column), false alarms (FA) when the participant reported a change on no-change trials (last row), and mislocalizations (ML), when participant incorrectly localized the change (all other off-diagonal elements).

##### Estimating psychometric and psychophysical parameters

Psychometric quantities (hit rates and false alarm rates) for each location were computed from the contingency tables. For orientation change detection with multiple change angles (n=5 participants), we also computed the psychometric function by fitting a three-parameter sigmoid curve to the hit rate as a function of change angle.

To estimate the psychophysical parameters (sensitivity and criteria), we employed a multidimensional signal detection model (the m-ADC model; Fig. 1D right), developed specifically for the analysis of behavioral data in multialternative detection tasks^24^. The model has been tested and validated widely for analyzing many kinds of detection tasks, both with humans and with non-human primates^25,38,39,42,43^. A detailed description of the model is available in these previous studies; we present a brief description here. In the 4-ADC model for our task the decision is modeled as multivariate Gaussian random variables. The decision variable distributions representing sensory evidence for change (signal) at each the four locations are represented along orthogonal axes. The decision variable distribution corresponding to no-change (noise) is centered at the origin. The mean of the signal distribution along each axis varies with the sensitivity (d’) for detecting change at that location. The decision boundary is parameterized by four decision thresholds (t), one for each location. On each trial, the observer indicates a change at the location at which the decision variable exceeded the corresponding threshold by the largest value. If the decision variable did not exceed threshold at any location, the observer indicates a no-change. We derive the criterion measure of choice bias as (c = t--d’/2)^25,44^. The criterion is inversely related to choice bias so that the lower the choice criterion at a location, the higher the choice bias towards that location. The m-ADC model was fit to response proportions in the stimulus-response contingency table, and sensitivity and criteria were computed using maximum likelihood estimation. Goodness-of-fit, estimated with a randomization test based on the chi-squared statistic, indicated that model fits did not deviate significantly from the data (p=0.6 [0.1-1.0], median [range]).

In the results in the main text, we compared psychophysical parameters at the ST location with the average values across the three SA locations; we also performed pairwise comparisons between the saccade-target ipsilateral and contralateral locations with a Bonferroni-Holm correction for multiple comparisons.

We also estimated the temporal dynamics of presaccadic attention’s effects on psychophysical parameters. For this analysis, we binned trials with similar intervals between final set onset and saccade onset, in non-overlapping bins of 50 ms duration. Psychophysical parameters were estimated by constructing contingency tables for each time window, with data pooled across participants. These temporal dynamics were then plotted aligned to saccade onset (250 ms before to 50 ms after); error bars were estimated with a jackknife procedure.

#### Orientation estimation task

For the orientation estimation task, we computed the orientation report error at the ST and SA locations. Performance was quantified with the mean absolute error (MAE): the absolute deviation (in degrees) of the reported orientation from the true orientation of the probed stimulus, averaged across trials. For the double set trials, we analyzed each block type – initial set probed, and final set probed – separately. For the single set and noise mask trials, we report results from only the initial set probed blocks, because the Gabor stimuli for these trials always occurred in the initial set. Essentially, for these trials we sought to avoid detrimental effects (e.g., due to mismatched temporal expectation) on orientation estimates of a Gabor stimulus presented in the initial set in a block in which the final set was consistently probed. Nonetheless, pooling behavioral responses from both block types yielded nearly identical results. We computed precision as the reciprocal of the non-parametric circular standard deviation^45^ of the mean-subtracted, signed estimation error distributions. In Figures 2B–E, for visualization, we also plotted the von Mises function with concentration parameter fit using maximum likelihood estimation (data pooled across participants).

To quantify the influence of the final set on the report of initial set (recency bias) and vice-versa (primacy bias), we computed “bias” for the probed stimulus relative to the biasing stimulus using the signed area under the curve^46,47^. Briefly, we binned trials according to the orientation of the biasing stimulus relative to the orientation of the probed stimulus (bin width=60°, sliding by 1°) and plotted the average estimation error in these trials for each bin, to obtain a response bias curve. We computed the signed area under this curve, in the range of 10° to 60° magnitude (positive and negative) of relative orientation of the biasing orientation to the probed orientation. A positive signed area indicates an attractive bias toward the biasing stimulus (higher area in the 1st and the 3rd quadrants), whereas a negative signed area indicates a repulsive bias away from the biasing stimulus (higher area in the 2nd and the 4th quadrants).

### Fitting behavior with the variable precision model

We modified the variable precision (VP) model to fit participants’ behavior in the multialternative change detection task. A detailed description of the standard VP model is available in van den Berg et al., 2012^26^; in our results, we call this the “baseline” model (Fig. 5A). Briefly, the VP model assumes that precision varies across items and trials. On each trial, the value of precision (*J*) at each location is drawn from a gamma distribution, parameterized by mean 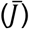 and scale (*τ*) parameters for that location, i.e., 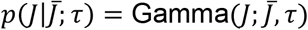. The internal measurement *x* of an external stimulus *θ* on each trial is assumed to follow a von Mises (VM) distribution, centered on *θ* and with a concentration parameter *κ* which determines the variance of the noise in measurement; the variance is inversely proportional to *κ*. The relationship between the concentration parameter, *κ* and precision, *J* is is denoted by *κ* = *φ*(*J*), and computed numerically. Therefore, the distribution of internal measurements is given by a mixture of VM distributions, with parameters drawn from the corresponding gamma distributions. Thus, the internal measurement is distributed as 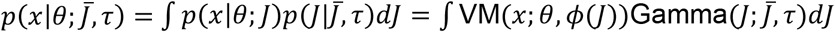. We approximated this distribution by averaging VM distributions corresponding to 1000 samples of *J* (drawn from the gamma distribution). To incorporate motor noise during the report 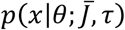 was convolved with another VM distribution with concentration parameter *κ_m_*; the latter parameter was fixed (*κ_m_*=25) based on its value in a previous study^26^.

#### Orientation estimation task

Based on evidence from our experiments, we modeled different mean precisions 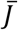 for the initial and final stimuli at the ST location (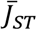 and 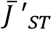, respectively), and a common mean precision at the SA locations 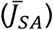. Along with these 3 parameters, we estimated a common *τ* (scale) across all locations. To ensure robust parameter estimates, we pooled the error distribution in the orientation estimation task across participants; this was done separately for the initial and final stimuli at the ST location but combined across stimuli at the SA locations. Model parameters were estimated with maximum likelihood estimation using grid-search by simultaneously fitting these three error distributions^26^.

#### Orientation change detection and localization task

Internal measurements corresponding to the orientations of the initial (***θ_i_***) and the final (***θ_f_***) stimuli are represented by latent vectors (***x*** and ***y***, respectively); these follow von Mises distributions with concentration parameters ***κ_x_*** and ***κ_y_***, respectively. For example, for each location *l* in the initial stimulus set, the internal measurement *x_l_* for each trial is drawn from a VM distribution with concentration parameter *κ_x,l_*, and centered at *θ_i,l_*. Similarly, for the final stimulus set, the internal measurement *y_l_* is drawn from a VM distribution with concentration *κ_y,l_* centered at *θ_f,l_*. Across trials, the concentration parameters vary depending on the values of precision *J* drawn from their respective gamma distributions. At the change location *c*, *θ_i,l_*=*c* and *θ_f,l_*=*c* differ by the change angle Δ*θ*. At the other locations, change angle was zero as per our actual experimental task design, so that *θ_i,l≠c_* = *θ_f,l≠c_*. The posterior odds ratio of change to no-change at each location 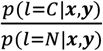 evaluates to a quantity proportional to 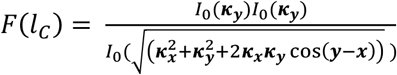, where *I_0_* is the modified Bessel function of the first kind of order zero^48^. To model change localization along with a no change response, we introduced a “decision threshold” parameter *η* in the model. If the posterior odds ratio does not exceed the decision threshold for any location *l*, the observer reports a “no change”. Otherwise, the observer reports a change at the location at which maximally exceeds the decision threshold at that location (argmax [*F*(*l_c_*) - *η*]).

We chose constraints on the model informed by the empirical results observed in the orientation estimation task. First, we modeled a common mean precision parameter for the initial stimulus at the ST location and the stimuli at the SA locations, because the mean precision estimates were comparable in the orientation estimation task paradigm. Second, because the mean precision for the final stimulus at the ST location was estimated to be six times the mean precision for the initial stimulus, we scaled the mean precision for the final stimulus at the ST location accordingly. Third, we used a common scale parameter, *τ*, across locations and stimuli, with value estimated from the orientation estimation task.

In the “baseline” model, the decision threshold, *η*, was fixed to 1. In the “perceptual bias” model, in addition, we modified the model to include a presaccadic recency bias *B* at the ST location. The internal measurement for the initial ST stimulus was modeled with a two-step process. First, an intermediate measurement *x_ST_* was computed with VM noise with *κ_x,ST_*. Next, to obtain the final measurement *x’_ST_*, we modeled a perceptual bias towards final ST stimulus (*y_i_*) as: *x’_ST_* = *B_y_ST__* + (1 - *B*)*x_ST_*, *B* ∈ [0,1]. In the “both biases” model, we additionally modeled differential thresholds at the ST and SA locations, which were estimated during model fitting. Finally, the “choice bias” model incorporated these differential thresholds as in the “both biases” model, but not a perceptual bias. Thus, the numbers of parameters for the baseline, perceptual bias, choice bias and both biases models were 1, 2, 3 and 4, respectively. The distribution of the MAP estimate, 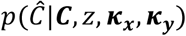, was computed through Monte Carlo simulation with 1000 samples of *J* and 4800 samples of ***x*** and ***y***.

Model fits were performed for each participant individually, with responses averaged across the SA locations. As before, model parameters were estimated with maximum likelihood estimation (multinomial likelihoods) using generalized pattern search. For formal model comparison, we employed the corrected Akaike Information Criterion (AICc)^49^, estimated independently for each participant. Goodness-of-fit was evaluated with a randomization test based on the chi-squared statistic. For 2/10 participants for whom the goodness-of-fit p-value suggested poor model fits (<0.05) for the both biases model, we re-ran the parameter estimation with an additional initial condition; this same initial condition was then tested across the choice bias model also. To avoid the potential for overfitting with the more complex models, we computed cross-validated data likelihoods using a 4-fold cross-validation using a “leave session out” approach. Each model’s predicted response proportions, obtained from the left out fold, were compared against observed response proportions using Pearson’s correlations.

### Statistical analyses

For change detection and localization tasks (both orientation and contrast), we employed a randomization test to test for significance of differences in sensitivity and criterion across the ST and SA locations (delta *d’* or delta *c*). Briefly, to estimate the *p*-value for a significance testing for sensitivity differences between these locations (*d’_ST_* – *d’_SA_*), we simulated response proportions for a “sensitivity-null” scenario by equating the sensitivity at all locations to the average *d’*, while retaining the criteria at their original values. With these sensitivity-null response proportions we generated 1000 contingency tables multinomial sampling while preserving row sums (number of trials per change event type). *d’*-s at each location were estimated from each of these contingency tables and a null distribution of delta *d’* was generated. The one-sided *p*-value represents the proportion of the values in the null distribution that exceeded the observed difference in *d’* between ST and SA locations. A similar approach was used to assess the significance of delta *c* (*c_ST_* – *c_SA_*) except that, in this case, a “criterion-null” scenario was simulated by equating the criteria at all locations to the average *c*, while retaining the *d’*-s at their original values.

For estimating significant differences in the analysis of sensitivity and criterion dynamics, we employed an identical randomization test, except that significance was evaluated independently in 7 non-overlapping 50 ms duration windows centered at −250, −200, −150, −100, −50, 0 and 50 ms with respect to saccade onset. Here, and elsewhere, unless stated otherwise, multiple comparisons correction was performed with the Bonferroni-Holm procedure. For orientation estimation, all the tests of significance for differences in estimation error, across the saccade toward and saccade away locations were performed using a permutation test comparing the observed differences against null distributions of differences. The null distributions were obtained either by using all possible permutations of location labels for comparisons with n=5 participants (single set and noise mask trials), or by random shuffling of location labels 1000 times for comparisons with n=10 participants (double set trials). For bias, we tested for significant difference from zero using a two-tailed Wilcoxon signed tank. Similarly, for comparing estimated model parameters across locations and conditions, we employed a non-parametric two-tailed Wilcoxon signed rank test.

Finally, we also computed Bayes factor (BF) based on the Jeffreys-Zellner-Siow (JZS) prior^50^ for pairwise t-tests (change detection tasks: one-tailed for criterion and sensitivity; orientation estimation tasks: one-tailed for estimation error and two-tailed for bias) in all pairwise comparisons between behavioral metrics at the ST and SA locations^51^. BF>3 (or >10) reflects substantial (or strong) evidence favoring the presence of the effect of interest. In addition, we examined how strength of evidence based on the BF changed as successive data points were collected, using Bayesian Sequential Analysis with the JASP software^52^. This analysis was performed for d’ and criterion for the visual change detection tasks, as well as precision for the initial and final stimuli, as well as the primacy and recency biases, for the orientation estimation task.

### Data and code availability

The data and custom scripts (Matlab and Python) for reproducing all figures in the paper are available at the following URL: https://dx.doi.org/10.6084/m9.figshare.21792002.

