## Supplementary Figures S1-S4 for "A surprising lack of presaccadic benefits during visual change detection"

### Supplementary figures and legends

**Supplementary Figure S1**

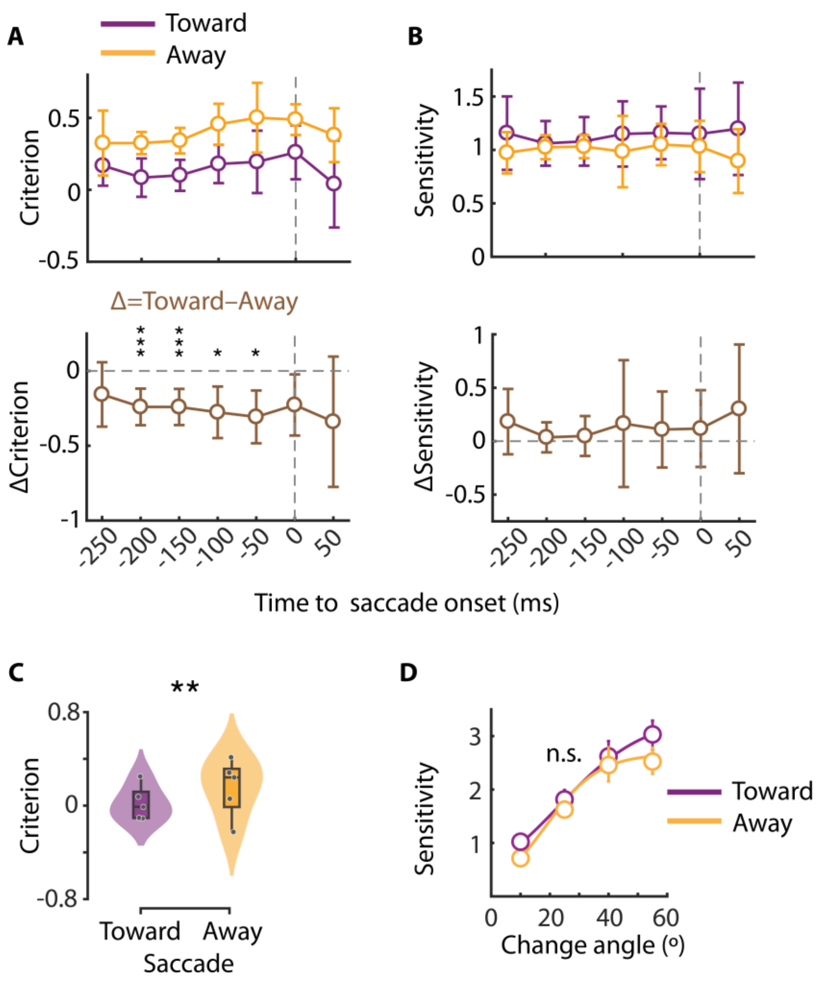


**Figure S1. Presaccadic attention effects in different time windows and for different stimulus strengths.**

**A.** Same as in Figure 1E, but showing temporal dynamics of criterion changes at the saccade Toward (purple) and Away (orange) locations. Other conventions are the same as in Figure 1E.

**B.** Same as in panel A, but showing temporal dynamics of sensitivity changes at the saccade Toward (purple) and Away (orange) locations. Other conventions are the same as in Figure 1E.

**C.** Same as in Figure 1E, but showing criterion at the saccade Toward and saccade Away locations (n=5 participants) for orientation change detection and localization with multiple change angles.

**D.** Psychometric function of sensitivity at the saccade toward and saccade away locations for orientation change detection and localization with multiple change angles (10°, 25°, 40°, 55°). Error bars: s.e.m. (n=5 participants).

**Supplementary Figure S2**

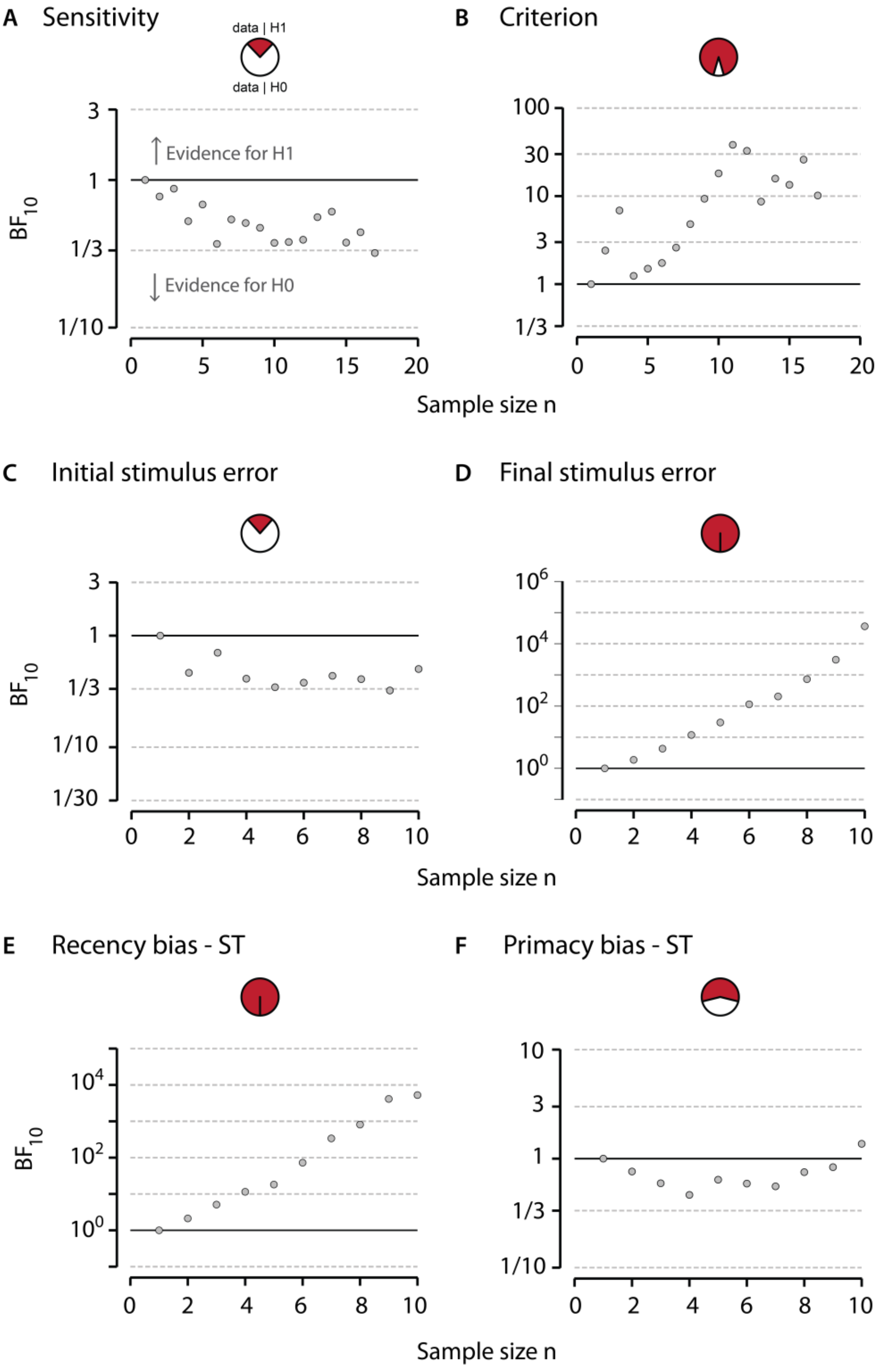


**Figure S2. Bayesian Sequential Analysis robustness check.**

**A**. Bayesian Sequential Analysis based on a paired sample t-test, as a function of sample size, for d' difference between the Saccade Toward (ST) and Saccade Away (SA) locations in the change detection tasks. Samples are pooled over the orientation change (n=10) and contrast change (n=7) detection experiments to yield a total sample size of n=17. Here, H0 is the hypothesis that d' is not different between the ST and SA locations while H1 is the alternative hypothesis that d' is different across these locations. The x-axis represents the sequential sample sizes (from n=1 to n=17 participants), the left y-axis indicates the Bayesian Factor supporting H1 over H0 ($BF_{10}$), and the right y-axis provides labels for different BF levels. Similarly, $BF_{01}$ quantifies the evidence supporting H0 over H1. (*Inset top-center*) Pie chart indicates the data likelihoods under the 2 hypotheses (white: H0 and red: H1).

**B**. Same as in panel (A) but for criterion difference between the ST and SA locations in the change detection tasks. Other conventions are the same as in panel A.

**C**. Same as in panel (A) but for a difference in initial stimulus precision between the ST and SA locations, in "double set" trials, in the orientation estimation task (n=10).

**D**. Same as in panel (C) but for a difference in final stimulus precision between the ST and SA locations, in "double set" trials, in the orientation estimation task (n=10).

**E**. Same as in panel (A) but for recency bias at the ST location in the orientation estimation task (n=10).

**F**. Same as in panel (E) but for primacy bias at the ST location in the orientation estimation task (n=10).

**B-F**. Other conventions are the same as in panel A.

**Supplementary Figure S3**

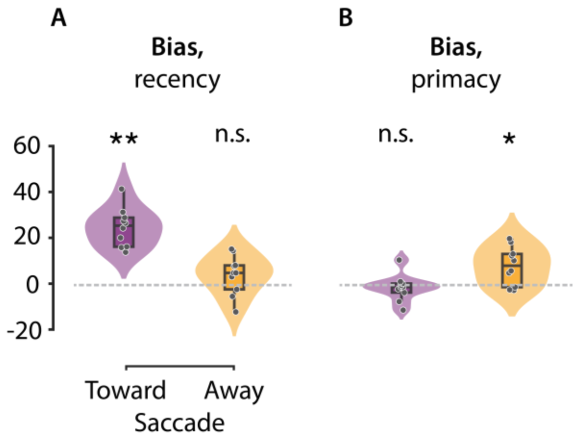


**Figure S3. Control analyses for temporal order reversal effects.**

**A.** Same as in Figure 3C (main text), but showing recency bias at the Saccade Toward and Saccade Away locations, following exclusion of trials in which the saccade onset occurred <50 ms after final stimulus offset.

**B**. Same as in panel A, but showing primacy bias at the Saccade Toward and Saccade Away locations.

(A-B) Other conventions are the same as in Figure 3C.

**Supplementary Figure S4**

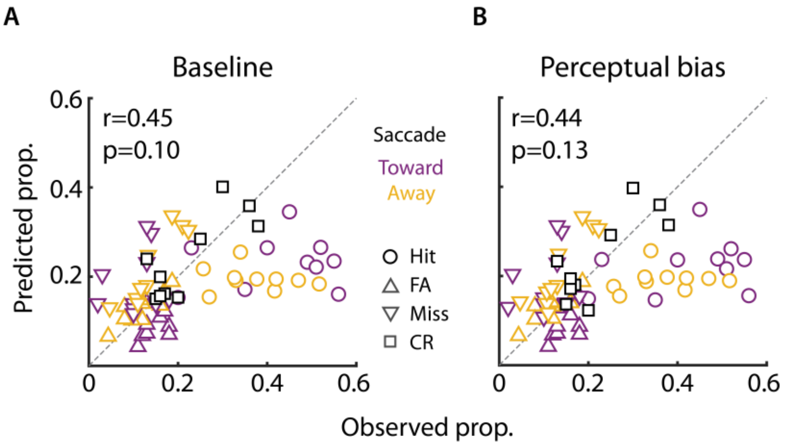


**Figure S4. Model predictions of behavioral responses in the change detection task.**

**A.** Same as in Figure 5E (main text), but showing response proportions predicted with the baseline model (y-axis) versus observed (true) response proportions (x-axis) in the change detection task.

**B.** Same as in Figure 5E (main text), but showing response proportions predicted with the perceptual bias model (y-axis) versus observed (true) response proportions (x-axis) in the change detection task.

**A-B.** Other conventions are the same as in Figure 5E.
